# Breaking the Tendon Repair Barrier: A Biomimetic Soluble Collagen Scaffold Enables Full Functional Regeneration of Achilles Tendon in Rabbits

**DOI:** 10.1101/2025.05.30.655895

**Authors:** Xiaoming He, Zhaohui Luo, Shenghua He

**Affiliations:** Independent researcher, Tokyo, Japan; Formerly: Wuhan Huayitongxin Biotechnology Co. Ltd, Wuhan, China

**Keywords:** biomimetic materials, soluble collagen, tissue engineering, regeneration, tendon

## Abstract

This groundbreaking research employs biomimetic artificial tendons for the first time globally to successfully regenerate a 2 cm defect in the rabbit Achilles tendon. The artificial tendon, woven by collagen threads derived from soluble collagen, is a biomimetic tendon with a composition, structure and function akin to natural tendons. The biomimetic tendon exhibits exceptional tensile strength of (43.4 MPa), surpassing that of the natural Achilles tendon (39.1MPa). The remarkable mechanical properties, coupled with its favorable biocompatibility and low immunogenicity, underscore the substantial clinical application potential of this biomimetic material.

A biomimetic tendon was implanted to repair a rabbit model of a 2 cm tendon defect without employing exogenous cells or growth factors. Twenty weeks after transplantation, the mechanical properties of the regenerated tendon were restored to approximately 80.1% of those of the natural Achilles tendon. This biomimetic tendon not only provides mechanical support during the initial phase of cell recruitment but also sustains strength to facilitate functional tendon reconstruction until material degradation and complete tendon regeneration. The regenerated tendon tissue exhibited regularly arranged dense collagen fibers along the longitudinal axis of the natural tendons, demonstrating high mechanical strength and seamless integration. Histological analysis further revealed a progressive enhancement in collagen fiber diameter, density, and structural integrity within regenerated tendons over time, closely resembling the characteristics of natural Achilles tendons eventually. Notably, there were no apparent immune reactions or inflammatory responses during the experimental cycle.

These findings unequivocally demonstrate the remarkable efficacy of biomimetic tendons in facilitating functional tissue regeneration, underscoring their significant medical and scientific value. This study not only presents a novel treatment approach for tendon injury repair but also establishes the foundation for future clinical tendon and ligament regeneration and repair, with extensive clinical application prospects.

## Introduction

Tendon tissue, characterized by its toughness and density, serves as a crucial conduit for transmitting tension between muscles and bones, facilitating bone and joint movement. In the realm of orthopedic medicine, tendon injuries are a prevalent occurrence. Various factors, including lesions, excessive load (such as physical exercise or lifting activities), strains, and others, can precipitate tendon damage, rupture, or even complete rupture. Consequently, these injuries can restrict joint stretching, impair motor function, and even compromise joint mobility. These adverse effects can lead to long-term dysfunction and a substantial decline in overall quality of life. Notably, some athletes have been compelled to end their careers due to tendon-related issues. Globally, tendon-related surgical procedures account for over 30 million annually, resulting in substantial medical expenditures of approximately 140 billion euros [Ganestam et al.,2016; Maffulli et al., 2011].

Compared with tissues such as skin or muscles, the regenerative ability of tendons is extremely poor. When the tendon is completely or partially ruptured, due to the characteristics of high tension, the two ends immediately retract after rupture, and it is difficult to dock naturally. Therefore, even under minor fractures, surgical intervention is required; it is almost impossible to heal itself under severe fractures or defects. This is due to that tendon is a dense connective tissue composed of highly ordered collagen fibers (mainly type I collagen). The number of tenocytes in a tendon is significantly lower compared to the number of cells in muscles or skin [Gross and Hoffmann, 2013; Tsiapalis et al., 2021; Qiu et al., 2014].

Traditional tendon repair methods primarily encompass direct suture, autologous tendon transplantation, and heterologous tendon transplantation. However, each method presents distinct limitations. Direct suture is restricted to small defects and carries a risk of poor healing due to excessive tension. Autologous transplantation offers benefits, but the donor tendon site is limited, the donor tendon lacks sufficient strength, and it damages the supply area and sacrificial function. Furthermore, the shape, size, and length of the donor tendon often do not align, particularly with underdeveloped tendons in minors, exacerbating the issue. Allogeneic tendons undergo repeated freeze-thawing, freeze-drying, radiation, chemical treatment, and other methods prior to transplantation. Nevertheless, due to the inability to completely eliminate allogeneic antigens and the destructive treatment methods, allogeneic tendons experience tissue aging and significant biological and mechanical property deterioration. Chemical residues can induce severe inflammatory reactions. Even with a weakened immune response after allogeneic tendon transplantation, there is often a high volume of inflammatory exudation and varying tissue metabolism levels, leading to allogeneic tendon dissolution and compromised healing performance, notably, allogeneic tendons pose a risk of transmitting diseases such as HIV, HBV, and HCV [5 Chen et al., 2009; Kannus, 2000; Liu et al., 2008_9_, Sensini,et al., 2018].

Decellularized animal tissues, such as pig dermis, fetal cowhide, and the submucosal layer of pig small intestine, cannot be directly utilized as artificial tendons due to antigenic concerns and limited tensile strength. Their primary application lies as an adjunct to tendon repair techniques[Zheng et al., 2005].

In the initial stages of research on artificial tendons, a substantial emphasis was placed on mechanical properties. Subsequently, the development of artificial ligaments, such as the Leeds-Keio and LARS systems, facilitated the incorporation of polyterephthalate (PET) synthetic fibers to emulate tendon fibers. Notably, researchers employed electrostatic spinning and melting electroptewriting technology to fabricate wavy, curved synthetic fibers, similar to the structure of natural tendons[Sensini,et al. 2018; Denver et al., 2012; Hochleitner et al., 2018; Onur et al., 2017]. Although synthetic fibers exhibit superior mechanical strength compared to natural tendons, their limited affinity for tendon cells presents a significant challenge in their cultivation and proliferation. Furthermore, the material exhibits poor integration capabilities with recipient tissue, potentially leading to inflammatory responses, scar tissue formation, and adhesion. Consequently, achieving genuine functional regeneration remains a substantial challenge.

In recent years, the advancement of tissue engineering and regenerative medicine has offered novel approaches to tendon repair. The synergistic combination of scaffold, cells, and biological active factors emulates the structural integrity and functional capabilities of natural tendons. Nevertheless, significant progress has not been achieved in this field [Youngstrom et al.,2016; Xu et al., 2013; Denver et al., 2012; Hochleitner et al., 2018].

The primary component of tendons is type I collagen, comprising approximately 70-80% of their dry weight. This highly organized parallel fiber bundle arrangement confers tendons with exceptional mechanical performance. This unique ultra-microstructure underpins the tendons’ remarkable mechanical properties [Kannus, 2000].

Natural polymer materials, including collagen, hyaluronic acid, laminin, and others, exhibit low antigenicity and high affinity, making them exceptional biomedical materials. However, none of these materials have been utilized in the fabrication of artificial tendons to date. Notably, soluble collagen, the primary component of tendons, has been successfully applied in forms of sponge and solution (gel) in the medical field. However, it exhibits weak mechanical strength, strong hydrophilicity, and high solubility, leading to rapid dissolution and degradation in vivo. Consequently, its further application in tissue engineering remains restricted.

The medical community urgently requires artificial tendons fabricated from high-strength, high-affinity, high-nutrient, and slow-degrading materials.

Given the aforementioned background, this research has ingeniously developed a technology that substantially enhances the strength of soluble collagen, decelerates its degradation rate, and constructs biomimetic tendons with low immunogenicity and exceptional mechanical strength, mimicking the characteristics of natural tendons. This biomimetic tendon not only provides initial mechanical support as a scaffold for cell migration, proliferation and differentiation, but also sustains a certain level of strength to facilitate the functional reconstruction of the host tendon until the material degradation, and the complete regeneration of its own tendon. The unequivocal regeneration of a 2 cm-rabbit tendon defect demonstrates the biomimetic tendon’s remarkable biocompatibility, degradation characteristics, and efficacy in promoting tendon regeneration. This research provides novel insights into the underlying mechanisms of tendon regeneration.

This report presents the world’s inaugural application of artificial materials to implement a comprehensive structural and functional regeneration in a large tendon defect model. This accomplishment not only establishes a novel benchmark within the field of tendon and ligament tissue engineering but also presents novel concepts and treatment strategies for patients with trauma, surgical defects, or congenital abnormalities.

## Materials and methods

### I. Artificial Tendon Production

#### 1.1 Collagen Thread Production

Dissolve a medical-grade collagen powder (collagen Type I) into a solution. Using a spinneret (e.g., a syringe needle), squeeze the collagen solution into a coagulation bath to form collagen threads. Subsequently, place the solidified collagen thread into a cross-linking solution to obtain a robust collagen thread. For further details, please refer to an article [He et al., 2025].

The collagen thread demonstrates exceptional mechanical strength, surpassing the standards of absorbable surgical sutures. It also possesses remarkable biological properties, as officially recognized by the Chinese Food and Drug Administration (CFDA) (No. MZ17010224, No. WT17080974, No. WT19010144, No. WT19010145). In animal experiments and preliminary human cosmetic plastic surgery, it is utilized as an absorbable surgical suture, demonstrating excellent therapeutic efficacy [He et al., 2025].

After immersing a collagen thread (monofilament) in physiological saline for an hour, it was subsequently compressed firmly on a steel plate. It was observed that the collagen thread is composed of numerous minute collagen fibers, similar to those found in natural tendon bundles as depicted in Figure 1. The collagen thread and tendon bundle exhibit remarkable similarities in structure, composition, and function (high strength).

**Figure 1.**
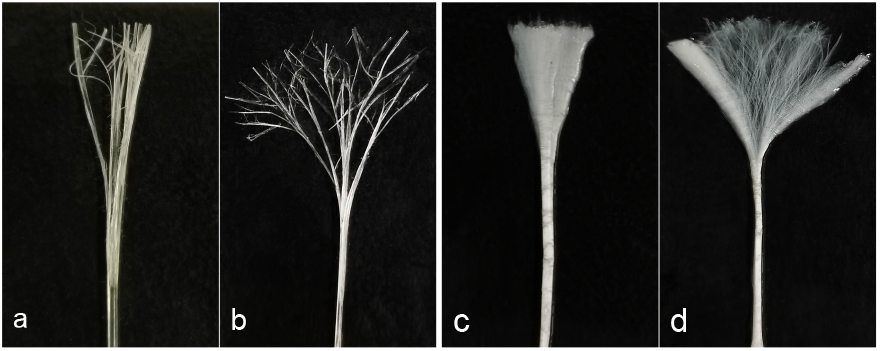
Structure comparison between a collagen thread and rabbit Achilles tendon. a,b: A collagen thread after being compressed; c,d: Rabbit autologous tendon bundle.

#### 1.2 Production of Artificial Tendons

Artificial tendons are fabricated from collagen threads through weaving (or twisting) techniques. The tensile strength of artificial tendons is influenced by two factors: the fabricating methods and the diameter of the threads. In this study, a diverse range of artificial tendons was produced using various techniques and thread diameters. Subsequently, their tensile strength was evaluated to identify the most robust artificial tendons.

Two methods are employed to fabricate artificial tendons using collagen thread with a diameter of 0.175±0.015mm. Subsequently, the tensile force in both dry and wet states is measured.

##### 1.2.1 Comparison of Fabricating Methods

###### Twisting Method

Multiple collagen monofilaments are twisted into a strand, and subsequently, multiple strands are twisted into artificial tendons (Figure 2-ab). This method involves interactive twisting (a) and co-twisting(b). The interactive twisting rope and the strand exhibit opposite twisting directions, enabling the internal residual stress and rotational moment to counteract each other. Consequently, the structure is compact and challenging to disperse, exhibiting resistance to bending. In this instance, the interactive twisting technology is employed; 12, 20 and 40 collagen threads are twisted to make artificial tendons respectively. The twisting parameters are shown in Figure 3-Table d.

**Figure 2.**
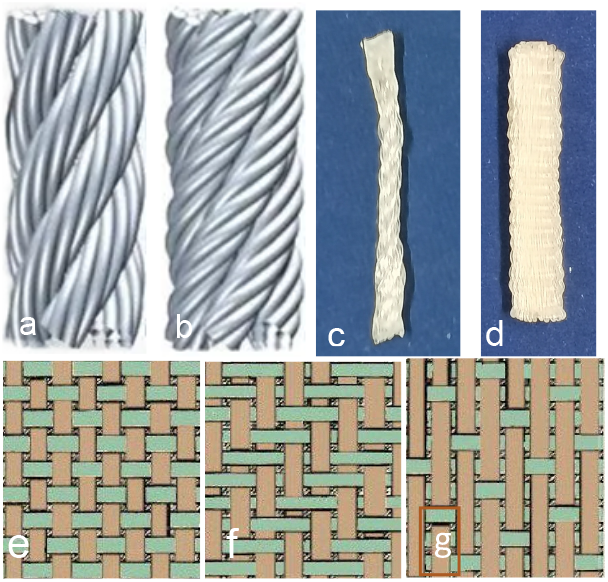
Fabricating methods of artificial tendon; a,b: twisting; c: Twisted tendon; e,f,g: weaving; d:weaved tendon.

**Figure 3.**
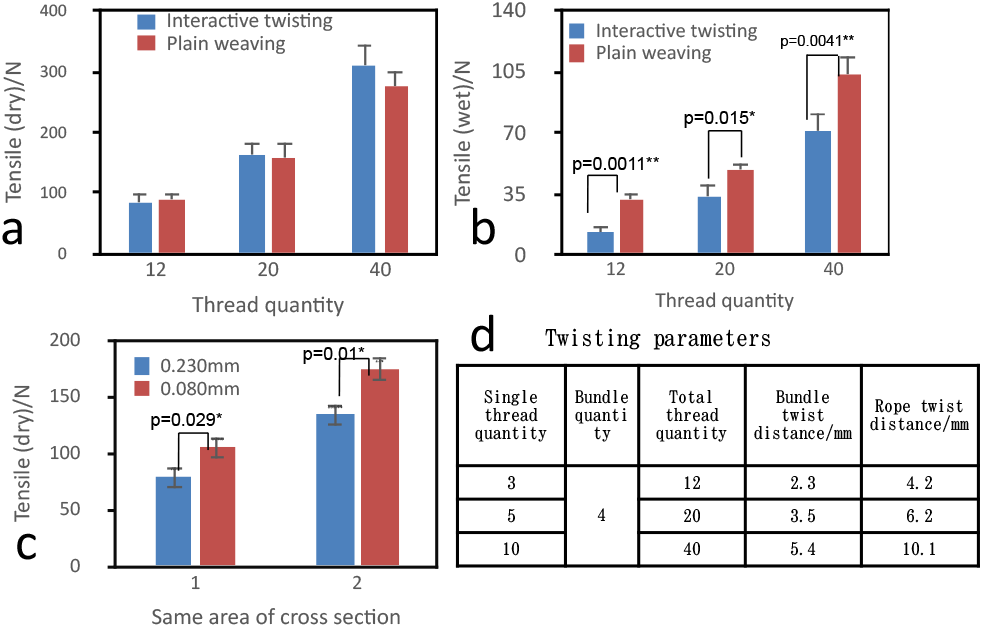
Screening among various fabricating methods. *p<o.o5, **p<0.01

###### Weaving Method

Collagen threads are woven into a sheet. The fabric is categorized into plain, twill, and satin structures (Figure 2efg). The plain structure exhibits enhanced tightness, resulting in high strength and wear resistance. In this case, plain weave is adopted, which is subsequently rolled into a cylindrical artificial tendon along the warp direction (Figure 2d).

Take the warp density of 95/10cm and the weft density of 500/10cm as the parameters on the weaving machine, and weave 12, 20 and 40 weft sheets to make artificial tendons.

###### Tensile Strength

Fix the two ends of the artificial tendon on the fixture of the mechanical testing machine, stretch at a uniform speed of 100±10mm/min until rupture, and record the maximum force (N). Immerse the artificial tendon in physiological saline for 1h to measure wet tensile strength, the results are shown in Figure 3-a (dry state) and Figure 3-b (wet state).

###### Conclusion

The t-test results indicate no statistical significance in the comparison of dry tensile forces. The p-values for p_12_=0.52, p_20_=0.76, and p_40_=0.08 are all greater than 0.05. Conversely, the comparison of wet fracture tensile forces reveals statistical differences, with p-values for p_12_=0.0011, p_20_=0.015, and p_40_=0.0041 being less than 0.05.

Considering the same number of collagen threads, the plain-woven artificial tendon demonstrates superior fracture and stretching capabilities, making it a more suitable option as an artificial tendon.

##### 1.2.2 Comparison of Collagen Thread Diameter

Artificial tendons are fabricated by utilizing collagen threads of varying diameters, while ensuring a consistent total cross-sectional area. Subsequently, their tensile forces are measured.

**Fabrication** using collagen yarns of 0.230±0.015mm and 0.080±0.010mm diameter

1. Collagen yarn with a diameter of 0.230 ± 0.015 mm is employed to fabricate sheets for the production of artificial tendons. The machine settings employed are as follows: warp density - 95/ 10 cm, weft density - 390/10 cm, and the number of weft yarns is 11 and 16.
2. Collagen yarn with a diameter of 0.080±0.010mm is employed to fabricate artificial tendons. The machine settings utilized are as follows: warp density -140/10cm, weft density -1140/10cm, and the numbers of weft yarns are 91 and 132.

###### Strength of Artificial Tendons

The strength was evaluated as described previously. The results are presented in Figure 3c.

###### Conclusion

A t-test was conducted for each of the two data groups. For the first group, p=0.029<0.05, indicating statistical significance. For the second group, p=0.010<0.05, also indicating statistical significance. These results suggest that the rupture stretching force of the artificial tendon fabricated from small threads is greater.

##### 1.2.3 Production of Artificial Tendon

In this study, artificial tendons were fabricated using a plain weave with a collagen thread diameter of 0.080mm. It was woven with the following parameters: warp density (140/10cm), weft density (1140/10cm), total warp roots (40), and number of threads per thread (4). The resulting textile obtained a 12mm×20mm sheet. Subsequently, the sheet was rolled into a cylinder along the warp axis (Figure 4a) and subsequently stitched with collagen threads. Finally, an artificial tendon with a length of 20mm and a diameter of 2mm was produced (Figure 4b).

**Figure 4.**
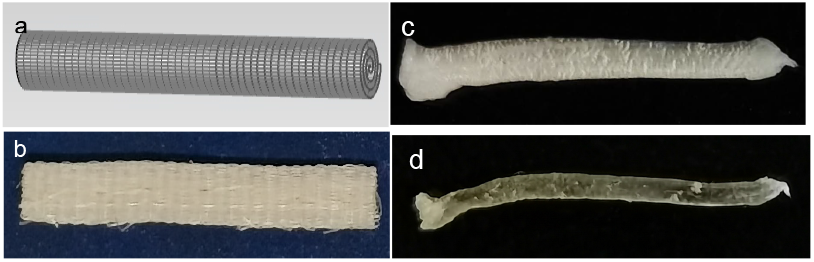
Artificial tendons and natural rabbit Achilles tendons a: Plain curl diagram; b: Artificial tendon for animal experiments; c: Fresh rabbit tendon; d: Dry rabbit tendon,

#### 1.3 Mechanical Strength of Artificial Tendons

##### 1.3.1 Tensile Strength

The tensile strength of dry and wet artificial tendons was measured, as depicted in Figure 5. The tensile strength of dry artificial tendons is substantially higher compared to that of autologous tendon. Notably, the tensile strength of dry artificial tendons exhibits a rapid decline upon immersion in physiological saline. There was no discernible difference in tensile strength between wet artificial tendons (43.4 ± 9.4 MPa) and rabbit Achilles tendon (39.1 ± 8.7 MPa).

**Figure 5.**
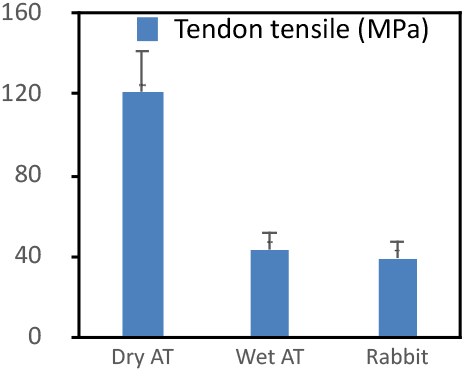
Comparative tensile strength of artificial tendon (AT) and rabbit tendon.

##### 1.3.2 Suture Strength

Suture strength is another crucial parameter of artificial tendons, which was measured as follows: thread the artificial tendon with 5-0 nylon sutures according to measurement method (Figure 6ab). Secure the ends of the nylon sutures together on the fixture on one side of the mechanical testing machine and stretch it uniformly at a speed of 100±10mm/min until the suture falls off. Record the highest strength. Soak the artificial tendon in physiological saline for 1 hour and measure the suture strength as above. The results are shown in Figure 6c. The suture strength of dry artificial tendons is substantially higher compared to that of autologous tendon, but there is no significant difference between the suture strength of the wet artificial tendon and that of the rabbit autologous Achilles tendon.

**Figure 6.**
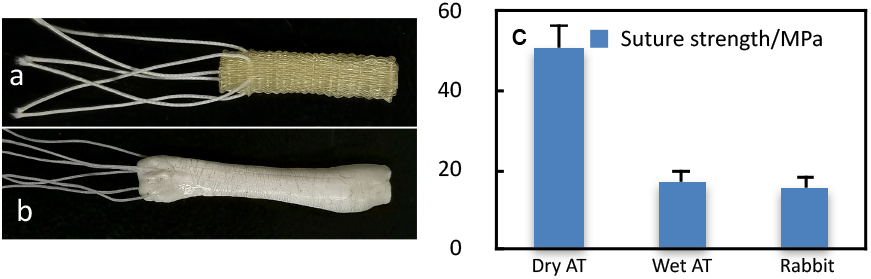
Suture strength of artificial tendon and rabbit autologous tendon. a,b: Methodology for determining suture trength; c: Comparison of rabbit tendon and artificial tendon (AT).

Subsequent animal experiments confirmed that the broken end of the tendon can be sutured securely. No transplanted tendon shedding was observed throughout the experimental cycle. Notably, the method of suturing tendons in surgery is different from that of suturing other tissues.

## 2. Animal Experiment: Repairing Tendon Defects

### 2.1. Animals

20 New Zealand white rabbits, ordinary class, male and female are not limited, weighing 20∼2.7kg, were from Hubei Experimental Animal Research Center.

All animal procedures were carried out in compliance with the applicable laws, regulations, and institutional guidelines regarding the use and care of laboratory animals. Animal welfare and ethical considerations were strictly observed.

‐ Housing: 12/12 light cycle, 22±1°C, 50% humidity;
‐ Food & drink: On-demand feeding;
‐ Approval: Ethics Committee of Wuhan University Central South Hospital (No. 2019007);
‐ Euthanasia: Anesthesia overdose.

### 2.2 Experimental Methods

General anesthesia is administered via a 10% hydrated chloraldehyde peritoneal injection at a rate of 1 mL/kg, combined with a fast-sleep-new II intramuscular injection at a rate of 0.3 mL/kg.

#### Tendon-Defect Model

Prepare the skin of the Achilles tendon in the hind limb. Make a midcutaneous incision on the posterior side of the Achilles tendon. Separate the subcutaneous tissue, expose the Achilles tendon and surrounding tissue, and cut open the membrane above the tendon. Separate the inner bundle (of the three tendon bundles) and retain the superficial outer and deep bundles. Create a tendon defect by removing approximately 2 cm of the middle segment of the Achilles tendon (Figure 7abc). Clean and preserve the excised tendon in physiological saline.

**Figure 7.**
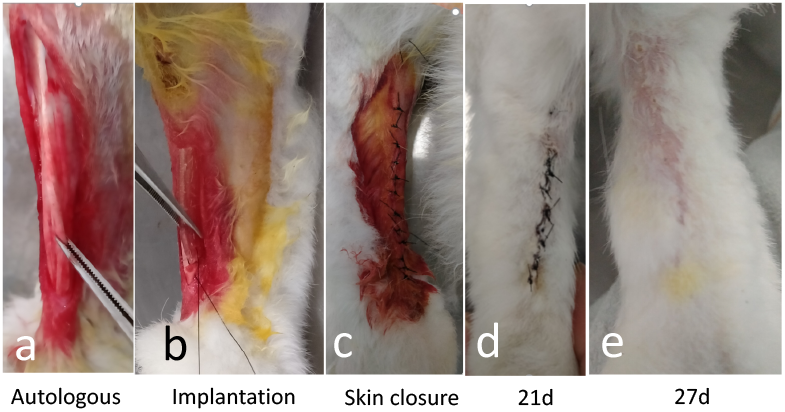
Transplantation of artificial tendon into the defect of Achilles tendon

#### Repairing Tendon Defect

Utilize a 5-0 nylon suture to connect the artificial tendon with the residual tendon of the rabbit model. Subsequently, close the tendon membrane with another 5-0 nylon suture. Finally, close the skin wound using a 3-0 silk suture. Administer penicillin daily twice for three consecutive days at a dose of 100,000 units each time.

### 2.3. Post-Surgery

Over time, the surgical site was excised to observe and document the surrounding tissue response, the healing of the anastomosis, and the regenerating tendon. Subsequently, the transplanted tendon was removed along with the newly formed tissue. The details are as follows:

Two rabbits underwent euthanasia due to excessive anesthesia administered at the second week post-surgery for a preliminary observation of inflammation, the stability of artificial tendons and tissue adhesion, etc. Three rabbits were euthanized at the fourth week post-surgery to further assess inflammation, immune response, wound healing, tissue adhesion, regenerative progress, degradation and the strength of artificial materials, etc.

Five rabbits were euthanized by excessive anesthesia at the 8th, 12th, and 20th weeks, respectively. In addition to the necessary observation, four samples were utilized to assess the strength and degradation rate of the transplanted tendons, while one case was utilized for histological analysis [Hematoxylin and eosin (HE) stain and Masson stain].

#### 3. Statistics

All data were presented as Mean **±** Standard Deviation (SD) and analyzed using Microsoft Excel software; t-test was employed to assess the disparity between the two groups; p-value less than 0.05 indicates a statistically significant difference.

## Results

### Appearance After Surgery

Following surgery (Figure 7abc), the experimental animals’ diets remained normal. The postoperative wounds exhibited slight redness and swelling, and occasional edema on the toes, which generally resolved within 3-5 days (Figure 7de). Throughout the experimental cycle, no apparent inflammatory reactions and immune responses were observed, nor did any transplanted tendons rupture or shed. Notably, one case of residual natural tendon degeneration, end-of-muscle fibrosis, loss of tendon elasticity, and less new tissue formation was identified.

### Regenerated Tendons

Two weeks after the implantation, the artificial tendons were removed and underwent evaluation. It was observed that a thin layer of newly formed white tissue covered the surface of the artificial tendon, approximately 0.04 mm in thickness. Upon separation by scalpel and scissors, the white loose tissue partially adhered to the surface of the regenerated tendon. At the 12th, 20th week after surgery, the white tissues were more evident (Figure 8d.d1.d2).

**Figure 8.**
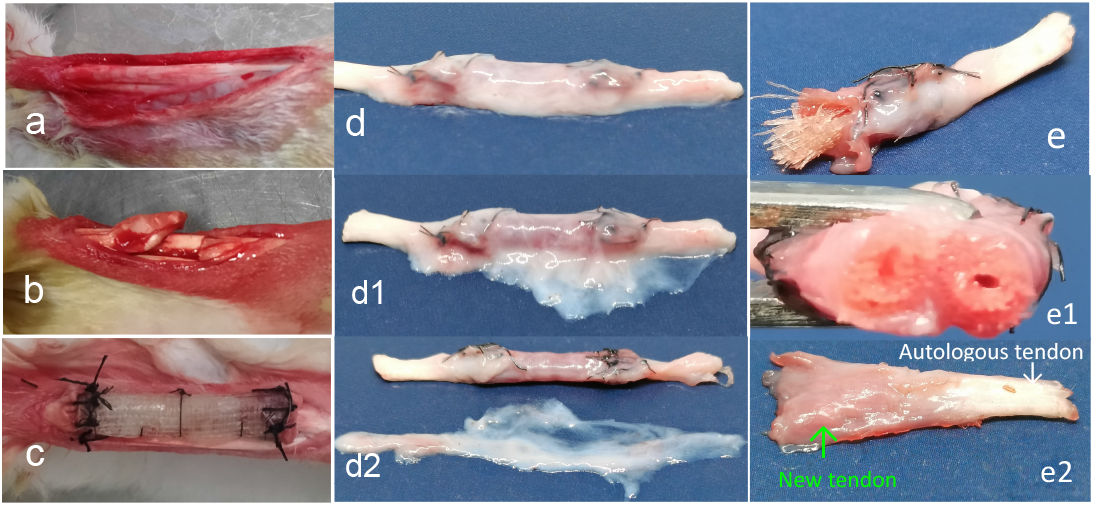
Images a,b,c: transplantation of artificial tendon; d, d1,d2: 20-week regenerated tendons with the white outer layer; d2: Separated outer white layer. e, e1, e2: Half of the 20-week tendon; e:Regenerated tendon with nylon sutures and collagen fibers; e1: Cross section; e2: Regenerated tendon without nylon sutures and collagen fibers; Green arrow: Regenerated tendon; White arrow: Autologous tendon. Right: Chronological changes in the thickness (mm) of the inner and outer layers of regenerated tendons.

At 4, 8, 12, and 20 weeks after surgery, the new tissue hierarchy on the implanted tendon exhibits a consistent pattern. It comprises a double-layer structure, with the white loose outer layer and the flesh-colored dense inner layer. This inner layer is connected to the residual end of the autologous tendon, while the outer layer is attached to both the inner layer and the autologous residual tendon (Figure 8 d,d1,d2).

Take out a portion of the outer layer of regenerated tissue and place it under a microscope for observation. At 4th, 8th, 12th, and 20th weeks after surgery, there is no discernible difference in the loose tissue of the outer layer. The outer layer contains a substantial amount of collagen fibers (Figure 9-abcd), the fiber morphology undergoes a gradual transformation from the outermost to the innermost regions. This transformation is characterized by the progressive transition of disorderly curving fibers into disorderly straight fibers, culminating in orderly straight fibers that align parallel to the longitudinal axis (Figure 9abcd).

**Figure 9.**
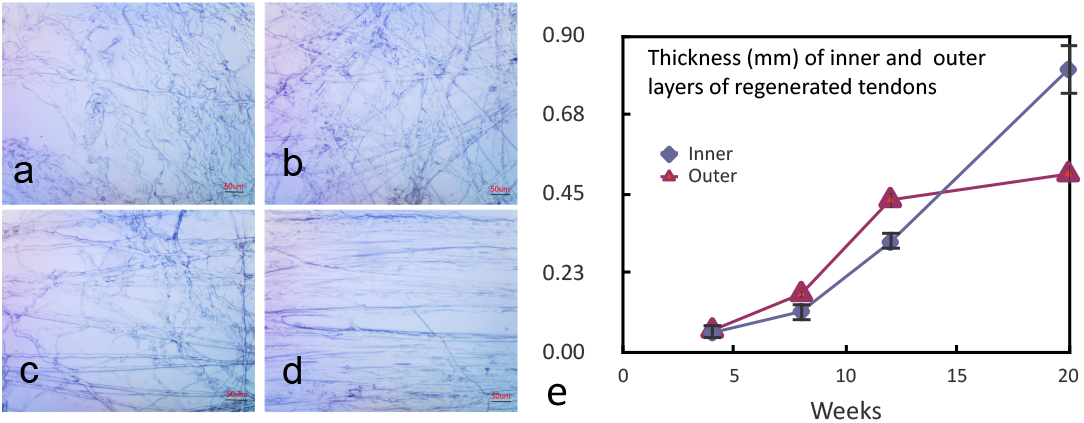
Left: Collagen fibers in the outer layer of regenerated tendon. Right: Chronological changes in the thickness (mm) of the inner and outer layers of regenerated tendons. abc: New tendons of 8,12, 20 weeks; a1b1c1: Disassembled new tendons of 8, 12, 20 weeks

The outer layer and inner layer undergoes thickness increase and transformation over time (Fig 9e), at first, outer layers is thicker than inner layer, and the thickness of both layers gradually increases, inner thickness surpasses that of the outer layer at 12 weeks. After 12 weeks, there is no indication of further thickening of the outer layer. The inner layer continues to thicken and acquire a tougher texture, resembling that of an autologous tendon. At 20 weeks, the inner layer becomes even tougher, and its thickness is much thicker than that of the outer layer.

Figure 10abc illustrates regenerated tendons after removing the white outer layer. The figure demonstrates that a inner layer of newly formed membrane envelops the artificial collagen tendon materials. This membrane is thin and can be easily separated from the artificial collagen tendon at an early stage, such as at 4 week postoperative. As the regenerated membrane thickens chronologically, it becomes increasingly challenging to separate it from the artificial collagen tendon at a later stage. By 20 weeks, the regenerated tissue has already integrated with the collagen tendon (Figure 10c1).

**Figure 10.**
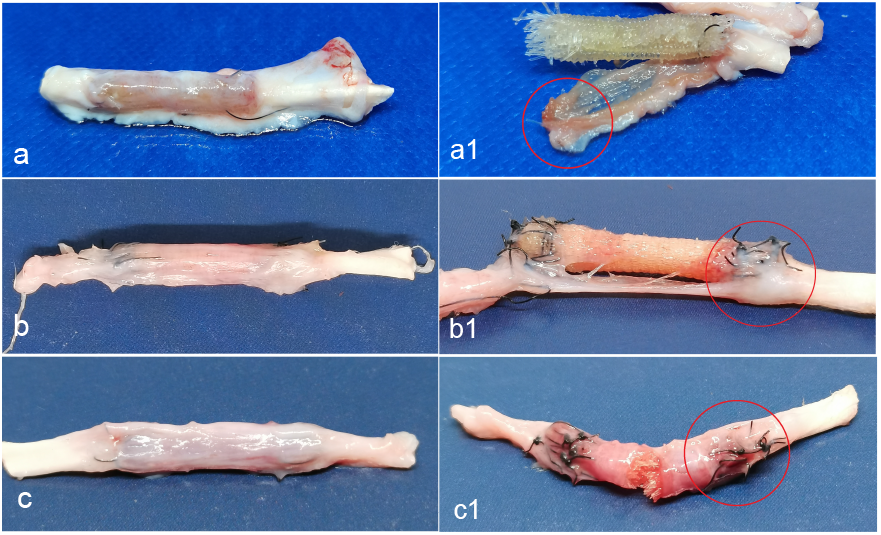
Regenerated tendons without white outer layer in chronological order. abc: New tendons of 8,12, 20 weeks; a1b1c1: Disassembled new tendons of 8, 12, 20 weeks

At 8 weeks post surgery, the inner layer exhibits a blood-colored (red) appearance (Figure 10a). Over time, it gradually transitions to a flesh-colored (pink) state at 12 weeks (Figure 10b). At 20 weeks, the inner layer develops a shallowly white appearance (Figure 10c), characterized by distinct strip-shaped areas that demonstrate enhanced resistance to deformation.

At 8 weeks, a distinct anastomosis exists between the implanted tendon and the autologous tendon (Figure 10a). Upon removal of the nylon sutures, the newly formed tissue and the residual autologous tendon are readily separated (Figure 10a1). The collagen fiber material is clearly distinct from surrounding tissue. However, at 12 weeks, the anastomosis boundary becomes indistinct, and the newly formed tissue has extensively infiltrated the artificial tendon, with the inner layer connecting to the residual tendon robustly. Even after suture removal, the anastomosis is challenging to tear and separate (Figure 10b1). Additionally, collagen fibers cannot be easily recognized because many cells and mucus have infiltrated the collagen fibers (Figure 10b1). At 20 weeks, a significant number of tissues have regenerated at the anastomosis site. Collagen fibers of artificial tendon cannot be recognized because they are wrapped by a thick regenerated tough tissue. Most of the collagen fibers in the artificial tendons have been degraded and absorbed, resulting in a seamless integration with surrounding tissue and autologous tendon. Even after removing the nylon sutures, it is difficult to separate the anastomosis site, and the connection becomes very firm and resilient (Figure 10c1).

### Histological Analysis (HE and Masson Stain)

Under the microscope, the rabbit’s autologous tendon (Figure 11N0) exhibits thick collagen fiber bundles arranged in a highly organized manner. There is a significant amount of matrix interspersed between the fiber bundles, and the bond between the fiber bundles is not particularly strong, allowing for easy disassembly.

**Figure 11.**
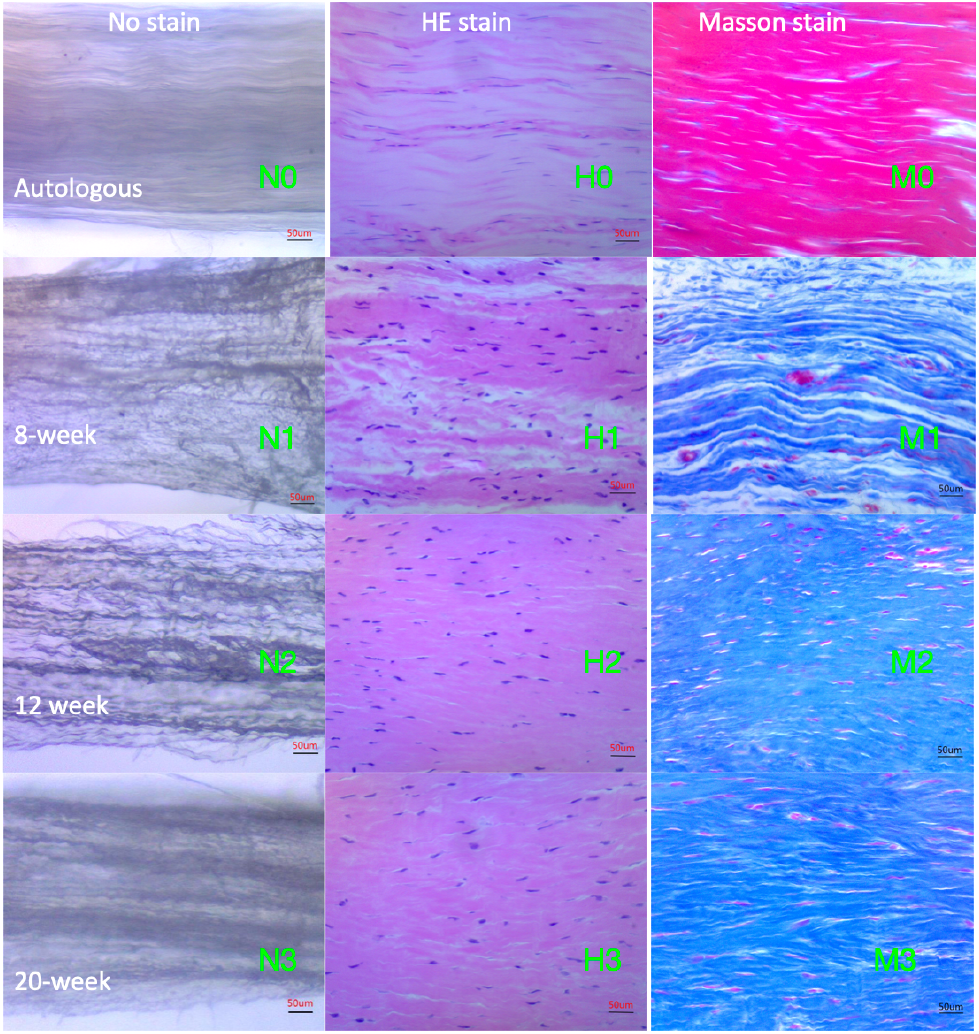
Microscopic structure of regenerated tendon and rabbit autologous tendon. Left column: Tissue without stain. Middle column: HE stain; Right column: Masson stain; Top first row: Autologous tendon; The 2nd row: 8-week regenerated tendon. The third row: 12-week regenerated tendon; Bottom row: 20-week tendon.

The microscopic structure of regenerated tendon were also observed in Figure 11. Over time, the increase in thickness and transformation of the collagen fibers of regenerated tendon becomes evident. At 8 weeks, ordered low-density tangled bonded collagen fiber bundles (small in diameter) are observed in Figure 11N1. At 12 weeks (Figure 11N2), the collagen fiber density increases, while the degree of fiber tangled bonding decreases. At 20 weeks, there is continuous increase in collagen fiber density, but the inter-fiber tangled bonding degree further diminishes(Figure 11N3),. The collagen fiber bundle of the autologous tendon can be easily separated without fiber rupture. Conversely, the adhesion of the regenerated tendon fibers is substantial, necessitating forceful separation and potentially causing occasional fiber rupture especially at initial stage, but it improved gradually over time and close to autologous tendon at 20 weeks.

Hematoxylin and eosin (HE) stains the nucleus blue and collagen fibers red. In contrast, Masson stains the collagen fibers of newly formed tendons blue and those of autologous tendons red.

HE stain of rabbit autologous tendon shows a large number of densely arranged thick collagen fibers (light dyeing) (Figure 11H0), and a small number of flat tenocytes between the fibers; new tendon tissue has low-density collagen fibers and a large number of oval tenocytes at 8 weeks (Figure 11H1); At 12 weeks, the collagen fiber density increased significantly, but it was still lower than the autologous tendon (Figure 11H2), and the number of tenocytes became low, cells became long oval; at 20 weeks, there was still increase in the density of collagen fibers, the number of tenocytes decreased further, and most cells became slender (Figure 11H3). Masson stain showed a similar results: a large densely arranged collagen fibers in autologous tendons (Figure 11M0); Masson staining can also observe that collagen fibers are getting denser and denser (Figure 11M1M2M3).

### Tensile Strength

The tensile strength of regenerated tendon was measured (Figure 12). The initial tensile strength of wet artificial tendon was relatively high, reaching 43.4 MPa, surpassing that of the autologous tendon. Subsequently, the tensile strength exhibited a notable reduction, with the lowest value at the 8th week, which was 25.1 MPa, representing 64.2% of the autologous tendon. Subsequently, the tendon began to rebound, reaching 31.3 MPa in the 20th week, closely approximating the autologous tendon’s value of 39.1 MPa, which represents 80.1% of the autologous tendon.

**Figure 12.**
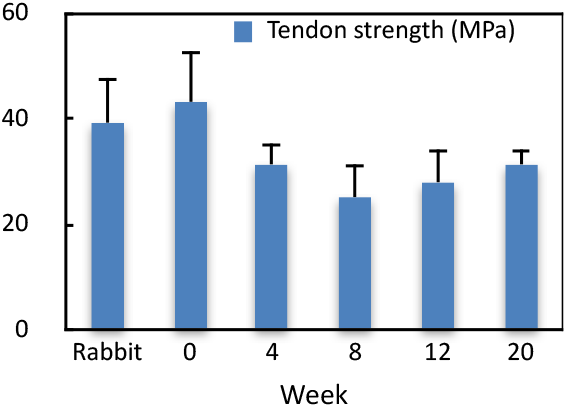
In vivo tensile strength of regenerated tendon postoperative. Rabbit indicates rabbit autologous tendon.

### Degradation of Implanted Tendons

Following the measurement of the tensile strength, the implanted tendon was dissected to extract the residual artificial collagen threads. These threads were subsequently washed and removed of any adhesive tissue. The collagen fibers were then dried and weighed. The degradation rate was subsequently calculated, as illustrated in Figure 13. It demonstrated a gradual increase in the degradation of artificial tendons over time, with a relative slowdown occurring after 12 weeks postoperative. At 20 weeks, the degradation rate reached approximately 62.1%, indicating a protracted period required for complete degradation. The observed decline in degradation rate from 12 weeks postoperative appears to be concurrent with the rebound in tensile strength. The tensile strength exhibited a rebound at 12 weeks (Figure 12). Figure 9 also demonstrated that the inner thickness is significantly greater than that of the outer layer from 12 weeks postoperative, suggesting that the augmentation speed of the number of functional cells, particularly tenocytes, has reached its peak at 12 weeks. As cells matured, they secreted a substantial volume of extracellular matrix (ECM) to augment the tensile strength of the regenerated tendon.

**Figure 13.**
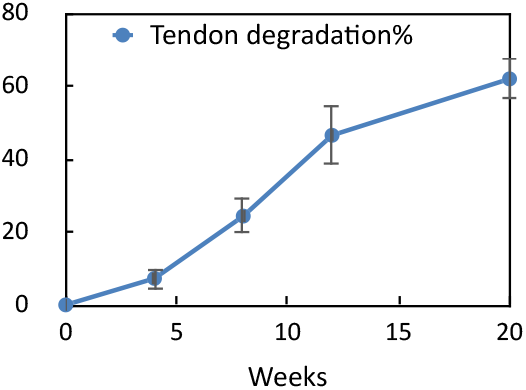
Degradation rate of artificial tendon in vivo chronologically

Simultaneously, as the secreted extracellular matrix (ECM, particularly collagen) accumulated and collagen fibers became increasingly dense, the alterations in the microenvironment would impact the physiological activity and metabolic performance of these functional cells, causing them to undergo transformations. The cell shape became slender, and even the number of tenocytes decreased. Consequently, the degradation of artificial tendon would progressively slow down at later stages. (Figure 13).

## Discussion

Tendon injury is a major challenge in sports medicine, orthopedics and tissue engineering. Regrettably, there is still no commercial artificial tendon to repair tendon defects in the medical practice so far.

This study fabricated a biomimetic artificial tendon that highly emulated the composition, structure, and function of biological natural tendons. The biomimetic tendons exhibited low immunogenicity, high mechanical strength, and biocompatibility. The experiment successfully demonstrated that biomimetic tendons facilitated tendon regeneration in the large defect model. It suggested that biomimetic collagen materials and their degraded products created a conducive environment for the growth and development of functional cells. Consequently, these cells differentiated and proliferated, ultimately promoting tendon tissue regeneration.

The three-tiered regulating mechanism, comprising a “collagen material, degradation product, and microenvironment,” appears to be a fundamental factor contributing to the successful regeneration of a substantial volume of tendon tissues.

### Existing Research and Medical Products

The regeneration capacity of natural tendons is significantly inferior to that of other tissues. Upon tendon rupture. The fibroblasts dominate the primary repair mechanism, resulting in the deposition of disordered collagen and the formation of scar-like tissue. This tissue exhibits structural and functional impairments, including diminished tensile strength, compromised elasticity, and heightened vulnerability to re-tearing. Consequently, tendon reconstruction is imperative.

However, it frequently encounters poor regeneration or even failure primarily due to antigenicity and the poor biocompatibility of transplant materials, allograft tendons and decellularized animal tendon matrices theoretically retain the original matrix structure and biological activity. However, potential immunogenic residues, such as membrane proteins (e.g., MHC I-type molecules), glycoproteins, or cell fragments, can trigger weak immune reactions and chronic inflammation, activating T cells and macrophages, secretion of pro-inflammatory factors (e.g., IL-1β, TNF-α, IFN-γ), and subsequent graft degradation or fibrosis hinder functional cells migration and recovery of function [Jiang at al., 2024; Voleti et al., 2012; Shi et al., 2019]. Furthermore, the graft exhibits inconsistencies with the host tissue in terms of metabolic rate, mechanical adaptability, and other characteristics, leading to stress concentration, local necrosis, and degeneration [Chen et al., 2009; Jiang at al., 2024; Voleti et al., 2012; Shi et al., 2019].

Currently, decellularized medical products such as Stryker-TissueMend, Conexa, and SIS (submucosal layer of the pig’s small intestine) are employed for tendon repair. However, the resulting “tendon-like tissue” exhibits limited mechanical capabilities and sliding functions, often accompanied by partial fibrosis or adhesion. Its mechanical properties are approximately half normal, and tissue maturity remains insufficient. This is because the decellularization process, such as using 1% TnBP and Triton X-100, cannot completely eliminate cell residues. This triggers mild immune and inflammatory reactions, leading to disordered collagen deposition (primarily type III collagen) and the formation of tendon-like tissue instead of functional tendons. Consequently, decellularized products are primarily utilized for small defects in shoulder sleeves and flexor tendons as an auxiliary role. They are not used to repair ligaments because joints are rich in immune cells, and synovial fluid amplifies inflammation and worsens scar formation. Additionally, poor blood supply and slow host cell migration and angiogenesis in joints hinder tendon regeneration.

Allograft tendons are better than decellularized animal matrix (Stryker: TissueMend, Conexa, SIS) but they are sourced from deceased individuals or abandoned limbs, posing risks of disease transmission and rejection [Shelton et al.,1997; Harner et al.,1996].

Zheng et al.[_38_2005] elucidated that biologically active materials facilitate functional tissue regeneration more effectively than inert synthetic materials. Martinello et al.[2014] demonstrated the application of self-assembled collagen gel in tendon repair, primarily focusing on small defect models and failing to substantiate its efficacy in large defect models.

Gross and Hoffmann [2013], Huang et al. [2023] reviewed various methods and materials for Achilles tendon defect repair in recent years and concluded significant challenges that remain, they claimed that tendon regeneration necessitates the integration of exceptional materials, intricate structures, exogenous cells, and growth factors.

Collagen materials can exhibit structural and functional similarities to natural tendons, prompting the synthesis of certain collagen materials through electrospinning and other advanced technologies. Nevertheless, the mechanical properties of these materials, such as tensile strength and elastic modulus, exhibit inferior characteristics in comparison to those of natural tendons. [Blackstone et al., 2021; Flores-Rojas et al., 2023].

Currently, numerous research endeavors in laboratories have been dedicated to the advancement of composite materials derived from collagen or other natural or synthetic polymers for tendon repair. Notably, some studies have explored the combination of collagen with other biologically active molecules, such as growth factors, to augment its capacity to facilitate regeneration [Wang et al., 2023; Rezvani et al., 2021]. Despite these numerous attempts, effective methods for repairing large tendon defects remain elusive.

This study has made substantial progress in the repair of severe tendon defects. Notably, this biomimetic tendon is predominantly fabricated using Type I soluble collagen, exhibiting exceptionally low antigenicity, high affinity, superior bio-integration, and remarkable mechanical strength. In contrast to allograft tendons, decellularized animal tendons, and other materials, this collagen biomimetic tendon presents several distinct advantages: its straightforward design and structure facilitate large-scale production and customization, while simultaneously ensuring safety and reliability.

### Mechanism of Biomimetic Collagen Tendon

This study elucidates the efficacy of biomimetic tendons in effectively bridging a 2 cm tendon defect in a rabbit model. The defect represents approximately 50% of the total length of the rabbit Achilles tendon, surpassing the size commonly defined as the “critical defect”[Müller et al., 2015; Tang et al., 2004]. Histological analysis reveals that the biomimetic tendon undergoes gradual invasion by host cells, resulting in the formation of a regular bundle of collagen fibers. This phenomenon can be attributed to the biomimetics to the natural tendon, including the arrangement of collagen fibers. This guide cell migration and differentiation in a specific direction [Sensini, 2018; Denver et al., 2012; Yin et al., 2010; Zheng et al., 2017; Angelina et al.,2018]. Notably, the regenerated tendon exhibits superior bio-mechanical properties, highlighting its clinical significance as an alternative to clinical tendon repair, thereby mitigating damage to the supply area. This aligns with the research findings of Martinello et al. and Zheng et al. which demonstrate the regenerative material’s ability to promote the differentiation of endogenous stem cells/ progenitor cells into tendon cells and supports the concept of “material-guided in vivo tissue engineering”.

Once the biomimetic collagen tendon establishes a connection with the natural tendon, revascularization and the infiltration of active cells initiate within the artificial tendon. Collagen scaffolds serve as a crucial mechanism in recruiting stem cells to the damaged site. They facilitate the migration of stem cells and guide their differentiation to specific cell types by establishing an optimal microenvironment [Engler et al., 2006; Guilak et al., 2009]. Collagen and its degradation products also play a crucial role in stimulating the formation of new blood vessels, which is essential for tissue repair and regeneration [Davis and Camarillo, 1995; Ruhrberg et al.,2002].

Fibroblasts, tenocytes, tendon derived stem cells (TDSCs), mesenchymal stem cells, and capillaries from the recipient invade and proliferate along the ends of the biomimetic tendon and between its collagen fibers. This process leads to the formation of numerous active cells, including tenocytes, stem cells, connective tissue cells, and capillaries. As these cells mature, they undergo a transformation into tendon tissue cells. These cells decompose and assimilate the coarse collagen of the biomimetic tendon, secreting fine collagen fibers that progressively replace the raw collagen fibers. This gradual formation of dense fiber bundles, neatly arranged along the longitudinal axis of the natural tendons, results in the transformation of the transplanted artificial tendon into a functional tendon tissue capable of metabolic activity.

This process closely resembles the transplantation of an allograft tendon [Harner et al., 1996; Shelton et al.,1997], which undergoes revascularization, cell proliferation, crawling replacement, collagen fiber reconstruction, and achieves the “resurrection” of the allograft. Notably, the regeneration effect of this artificial biomimetic tendon seems to surpass that of allograft tendons, potentially attributed to the absence of immune response and reduced inflammation.

### Postoperative Strength of Biomimetic Tendon

The biomimetic tendon’s strength before and after transplantation is a crucial determinant of its success. Eight weeks after transplantation of this artificial tendon, a significant reduction in mechanical strength was observed, which was subsequently maintained and gradually recovered. Although the lowest value (25.1 MPa) corresponds to 64.2% of the autologous tendon, it can sustain fundamental activity until it is fully restored. This is because the collagen materials of the artificial tendon undergo enzymatic degradation (surface erosion), and cell-secreted collagenases, such as MMP-1, MMP-8, and MMP-13, specifically recognize and degrade collagen [Pittayapruek et al., 2016]. This degradation process is more controllable, preserving structural integrity and mechanical properties. The degradation products of collagen are peptides and amino acids. These compounds play a multifaceted role in organisms, including intercellular communication, the regeneration and repair of signal transmission from the cellular to the tissue level [Shoulders and Rainers, 2009; Ricard-Blum 2011]. Consequently, collagen holds a crucial “nutritional” value for functional cells. As a natural component of the organism, collagen and its byproducts exhibit superior biocompatibility, facilitating cell adhesion, proliferation, and migration. During the degradation process, new intermolecular cross-linking or recombination may occur, temporarily strengthening the remaining structure [Hematti, 2018]. Additionally, collagen’s enhanced integration with host tissue promotes the growth of tissue cells on the suture and the substantial secretion of new extracellular matrix (ECM), especially collagen, which also help to maintain its strength [Brown and Philips, 2007]. When a large number of collagen fibers are densely arranged neatly along the longitudinal axis of the natural tendons, regenerated tendon has functioned normally and has restored the normal mechanical strength.

The mechanical strength decline in allograft tendon transplantation is more pronounced. Partial reasons are inflammation and immune response triggered by some components in the allograft, T cells and macrophages are activated to secret pro-inflammatory factors (e.g., IL-1β, TNF-α, IFN-γ), and subsequent resulting in graft degradation [Jiang at al., 2024; Voleti et al., 2012; Shi et al., 2019].

Notably, following the transplantation of this artificial tendon, the collagen fibers of the regenerated tendon, particularly during their initial phases, exhibit small, sparse, and irregular arrangements. Consequently, the initial regenerated tendon tissue possesses a lower strength compared to that of the autologous tendon. However, as the regeneration process progresses, the number of functional cells, including TDSCs, MSCs, tenocytes, increases, and collagen fiber secretion becomes more abundant. The fiber arrangement also becomes more regular, resulting in an enhancement of the strength of the regenerated tendon. This progression, as outlined by Shadwick [1990], elucidates the changes in mechanical strength of regenerative tendons and simultaneously elucidates the mechanism of regeneration.

Collagen materials possess the ability to retain their strength for a specific duration even within an active metabolic environment in vivo. This characteristic distinguishes them from synthetic polymer (such as PGA, PLA, and PLGA). Synthetic polymers are primarily degraded through hydrolysis in vivo, a process that is inherently uncontrollable. Consequently, this degradation leads to a progressive deterioration of structural integrity and mechanical properties, resulting in a rapid decline in mechanical strength [He et al, 2025].

### Tissue Adhesion

A notable challenge in this study lies in the pronounced adhesion of the regenerated tendon to the surrounding tissue. This restriction hinders the tendon’s sliding and stretching capabilities, adversely impacting its overall functional recovery. In contrast to the transplantation of autologous tendons, the biomimetic tendon scaffold is devoid of endogenous cells and does not undergo seeding of exogenous cells prior to transplantation. Consequently, this type of healing process would lead to the formation of adhesions.

Tendon adhesion is a prevalent complication following tendon repair, even in transplantation employing an autologous tendon. Usually, it can be mitigated through surgical intervention within the initial half-year. However, in this study, severe adhesion is observed due to the absence of endogenous healing mechanisms in artificial tendons, which are devoid of living cells. Consequently, exogenous processes of tendon regeneration lead to severe adhesion.

Specifically, the fibroblasts, mesenchymal stem cells, tenocytes and capillaries surrounding the tendon (synovial membrane, sheath, peri-tendon, etc.) invade the anastomosis end and proliferate, secreting collagen to facilitate repair. This process would results in adhesion, which appears to be an inevitable and crucial link in tendon healing.

In this study, after an artificial tendon was transplanted to repair the defect, the artificial tendon’s surface was covered by peri-tendon tissue, which relies on blood circulation for nutrient supply. The growth of surrounding tissue cells and the development of new blood vessels are the sole means of achieving successful regeneration, resulting in a gradual and adhesive healing process.

Adhesion is also associated with inflammation, scar formation, and the destruction of the sliding interface surrounding the tendon [Legrand et al., 2017]. Surgical trauma and suturing can cause tendon splitting, which is another significant factor in inducing adhesion [Wong et al., 2009]. To address adhesion, potential solutions include: (1).Seeding a large number of autologous stem cells (or tenocytes) before transplantation to enhance endogenous healing. (2).Supplementing bFGF and Vitamin C, which has been reported to increase tenocytes in the tendon and decrease adhesion [Caliari, and Harley, 2011; Omeroğlu et al., 2009]. (3).Utilizing TGF-β neutralizing antibodies to improve tendon activity [Li and Luo 2023] (4).Applying anti-adhesive substances, such as hyaluronic acid or polyethylene glycol derivatives, or anti-inflammatory drugs, on the surface of the biomimetic tendon to establish a physical barrier [Ishiyama et al.,2010]. (5).Integrating a sustained-release system of anti-fibrosis drugs (such as 5-fluorouracil or septhromycin C) into the artificial tendon [Zhao et al, 2015]. Additionally, exocrine-based cell-free therapy shows potential in reducing inflammation and promoting tissue repair [Shen et al., 2016], which warrants further exploration.

Repairing the paratendon tissue can reduce adhesion by protecting the blood supply of the tendon to facilitate endogenous healing. the paratendon tissue barrier also prevents the growth of granules [Lee and Fun, 2024]. Adhesion is also related to exercise after transplantation. Exercise at an early stage is beneficial for tendon sliding, which also facilitates collagen synthesis and reconstruction along the mechanical direction, as well as the longitudinal arrangement of new blood vessels, promoting substance exchange between tendons and humor fluids. This accelerates regeneration, repair, and shaping, effectively alleviating adhesion [Tam and Baar, 2025].

In the early stage, proper exercise should be protected by braces to prevent the biomimetic tendon from cracking.

### Research Limitations and Future Work

This study yielded promising results, but several limitations need to be addressed in this study. (1).Model Suitability: The present study is confined to the rabbit model. The artificial tendon should be further explored for tendon repair in large animals, including sheep, pigs, and humans. (2).Sports Conditions: rabbits were kept in a small cage, it is necessary to increase their exercise to improve the tendon repair. (3).Regeneration of Tendon-Muscle and Tendon-Bone Interfaces: In-depth analysis of the regeneration of tendon-muscle and tendon-bone interfaces is lacking. (4).Dynamic Balance Mechanism: The study fails to elucidate the detailed dynamic balance mechanism of material degradation and tissue regeneration. (5).Tendon Regeneration and Observation Period: The tendon regeneration period is insufficient, necessitating an extended observation period to evaluate the long-term stability of the regenerated tendon.(6).Lack of Molecular Mechanisms and Signaling Pathways: The study primarily focuses on histological and bio-mechanical evaluation, neglecting the crucial role of molecular mechanisms and signaling pathways in understanding the regeneration process. Future research should prioritize the analysis of relevant gene expression and proteome to gain a more comprehensive understanding of the biological characteristics of regenerative tendons.

Another concern is the protracted tendon regeneration process. The complete degradation of artificial tendons appears to require at least one year, and the complete restoration of mechanical strength and performance of regenerated tendon may take longer than one year. Consequently, the artificial tendon materials may still require refinement to facilitate rapid cell invasion and regeneration. as Sensini et al.[2017] demonstrated. The slow tendon regeneration can be attributed to the following reasons:

Firstly, blood supply to the tendon itself is compromised, particularly in the central region. Consequently, vascular regeneration lags, leading to inadequate nutrition and the inability to expel metabolic waste. These conditions impede cell activities. Secondly, it is reported that nerve tissue can stimulate tendon/muscle growth [Bean et al., 2023]. However, the regeneration of nerve may not have kept pace with tendon regeneration in the study. Thirdly, the distinctive characteristics of regenerated tendons at later stage, including their high collagen fiber density and relatively low cell density, may contribute to a gradually slow regeneration process and relatively weak strength [Kato et al., 1989].

In response to this, future researches can explore the following improvement strategies: (1) pre-culture stem cells, fibroblasts or tenocytes in biomimetic tendons and then implement implantation; (2) integrate angiopoietic factors (such as VEGF) in biomimetic tendons to promote neovascular formation [Lai, et al., 2022]; (3) imitate nerve function by providing electrical stimulation; (4) combine growth factors such as BMP-12, GDF-5, or IGF-1, which have been proven to promote the proliferation and differentiation of tendon cells [Ruiz-Alonso 2021]; and (5) optimize material porosity and connectivity to improve cell migration conditions and nutrient exchange to accelerate the regeneration process [Chen et al., 2009; Lomas et al., 2015; Xu et al.,2013].

### Clinical Application

Despite the aforementioned limitations, this study provides valuable insights and potential solutions for clinical tendon repair. Biomimetic tendons offer distinct advantages in addressing large defects, providing novel options for intricate cases that are challenging to manage using conventional methods. Notably, this biomimetic material holds special clinical significance for patients with limited options for autologous transplantation, such as elderly individuals or those with multiple injuries. The structural and functional similarities between tendons and ligaments enable the utilization of this biomimetic material as an artificial ligament, which is a highly anticipated product for numerous patients who rely on synthetic polymer materials like PET.

As a “ready-to-use” medical material that does not necessitate exogenous cells, growth factors, and can be produced on a large scale, its low cost and broad clinical adaptability are anticipated to transform into a groundbreaking product in the field of regenerative medicine.

Collagen, the fundamental skeletal protein, also found in vessels, esophagus, trachea, and other tissues, similarly, this material technology offers innovative solutions for addressing defects of these soft tissues by creating artificial vessels (especially small vessels), artificial esophagi, and artificial tracheas etc..

This technology holds a broad and promising future, with enhanced application prospects when combined with 3D printing and stem cell technologies.

## Conclusion

This research has successfully developed a biomimetic tendon with soluble collagen as its primary component. The tendon has demonstrated remarkable efficacy in repairing extensive defects in the rabbit Achilles tendon. Experimental evidence indicates that this material exhibits low immunogenicity and favorable mechanical properties. Although the regeneration process is relatively slow and tissue adhesion persists, the final regenerated tendon exhibits comparable structural and functional characteristics to the autologous tendon. These findings offer novel concepts and potential solutions to address the challenge of repairing severe tendon defects in clinical practice.

